# Recurrent acquisition of alien ribotoxin encoding genes by insect genomes

**DOI:** 10.1101/2019.12.20.880922

**Authors:** Walter J. Lapadula, María L. Mascotti, Maximiliano Juri Ayub

## Abstract

Ribosome inactivating proteins (RIPs) are RNA *N*-glycosidases that depurinate a specific adenine residue in the conserved sarcin/ricin loop of 28S rRNA. These enzymes are widely distributed among plants and bacteria. Recently, we have described RIP genes in mosquitoes belonging to the Culicinae subfamily (*Aedini* and *Culicini tribes*). We have also shown that these genes are derived from a single event of horizontal gene transfer (HGT) from a prokaryotic donor. In the present work, we show the existence of two RIP encoding genes in the genome of the whitefly *Bemisia tabaci* (a hemiptera species belonging to the Aleyrodidae family, distantly related to mosquitoes). Contamination artifacts were ruled out analyzing three independent *Bemisia tabaci* genome databases. In contrast to mosquitoes RIPs, the whitefly genes harbor introns and, according to transcriptomic evidence, are transcribed and spliced. Interestingly, phylogenetic inference combined with taxonomic distribution strongly supports that whitefly RIP genes are derived from an independent HGT event from a plant source. Our results suggest that RIP genes fill a functional niche in insects.

## INTRODUCTION

Horizontal gene transfer (HGT) is the reproduction-independent transmission of genetic material between organisms of different species. HGT has been reported to have occurred in all the three domains of life and is accepted as an important evolutionary force in prokaryotes [1-3]. On the contrary, its impact on multicellular eukaryotes (*e.g.* metazoans) is largely controversial [4]. In arthropods, many well-supported HGT events from bacterial or fungal sources have been described [5, 6]. Moreover, it has been suggested that HGT played a role in the herbivory of several arthropods and nematodes (see [7, 8] for a review)

Ribosome inactivating proteins (RIPs, EC 3.2.2.22) are RNA *N*-gycosidases depurinating ribosomes in the conserved alpha-sarcin/ricin loop of 28S rRNA, leading to irreversible arresting of protein synthesis [9-11]. RIP encoding genes are widely distributed in plants and scarcely within bacterial and fungal lineages [12]. Previously, we demonstrated the presence of RIP genes in genomes of mosquitoes belonging to the subfamily Culicinae [13]. This was the first description of the existence of RIPs in metazoans. Our research also evidenced that these genes are derived from a single HGT event from a bacterial donor species [14]. Recently, evidence has been accumulated on the role of *Spiroplasma* spp RIPs in the protective mechanisms generated by these endosymbiotic species against arthropod infection by natural enemies (see [15] for a review).

In this piece of research, we demonstrate that RIP genes are also present in the genome of a second lineage of insects: the hemiptera *B.tabaci*. Notably, these genes are embedded into a plant RIPs clade and are phylogenetically distant from mosquitos’ homologues. The collected evidences are consistent with an independent acquisition event of a RIP encoding gene by insects. Our results suggest that RIP genes may fill an important functional niche in insects leading to the recurrent selection either from horizontally transferred genetic material or by a symbiotic interaction.

## METHODS

### Homology search and sequence analyses

BLASTp and TBLASTN homology searches were performed under default parameters on metazoan databases using as query a previously reported set of RIP sequences [14]. Two automatically annotated protein sequences (GenBank: XP_018902206, XP_018902288) were retrieved from the *B. tabaci* MEAM 1 genome database. Pfam analysis was performed to confirm the presence of RIP domain (PF00161) [16]. Homology models were generated using Swiss-Model server [17] and visualized in PyMOL. Active site residues [18] were detected by sequence alignment to momordin (PDB 3my6).

### Genomic context analyses

The contig region containing RIP genes from *B. tabaci* MEAM 1 (NW017547285) [19] was compared employing Mauve software [20] to the corresponding genomic fragments of other members of the *B. tabaci* complex; namely *B. tabaci* SSA-ECA (PGTP01000858) [21] and *B. tabaci* MED/Q (ML134445) [22].

### Transcriptomic data analysis

The full RNA-seq dataset expressed as Reads Per Kilobase of transcript per Million mapped reads (RPKM) of *B. tabaci* MEAM 1 was kindly provided by Dr. Zhangjun Fei [19]. The logarithm of average level (among different experimental conditions) of gene expression for BtRIP1 (Gene ID: Bta13094) and BtRIP2 (Gene ID: Bta13103) were plotted along with boxes and whiskers graphs showing quartiles for the whole set of *B. tabaci* genes using GraphPad Prism version 5.00 for Windows.

### Multiple sequence slignment and phylogenetic inferences

The new *B. tabaci* RIP protein sequences were incorporated to our previous dataset [13]. The conserved region in the RIP domain (from Y14 to S196 residues, according to trichosanthin; number GenBank AAT91090) was selected for alignment, as it was reported [13, 14, 23]. MSA was constructed using MAFFT 7 online server [24] employing BLOSUM 30 as scoring matrix. Poorly aligned regions were trimmed as blocks. This MSA was used to perform the phylogenetic analysis by Maximum Likelihood method in RAxML 8.2.12 on XSEDE [25]. The WAG substitution matrix was selected as best model and gamma distribution with invariable sites parameter calculated using ProtTest3.4 [26]. To estimate the robustness of the phylogenetic inference, 100 rapid bootstrapping (BS) was selected. Transfer bootstrap expectation was applied to the obtained BS values in BOOSTER [27]. Phylogenetic relationships and divergence times among species from different taxa were obtained from Time Tree knowledge base [28]. FigTree 1.4.2 was used to visualize and edit the trees.

## RESULTS AND DISCUSSION

### 1. *Bemisia tabaci* genome harbors two RIP encoding genes

In the continuous quest for new RIP genes across the tree of life, we found two RIP encoding sequences in the hemiptera *B. tabaci* MEAM 1 (hence forward named BtRIP1 and BtRIP2). **Figure 1A** shows a schematic representation of the genomic scaffold (NW_017547285) harboring these genes. In contrast to RIP genes from mosquitoes, BtRIP1 and BtRIP2 genes contain one and two introns, respectively. This rules out the chance of bacterial contamination, a frequent artifact of massive sequencing projects [29]. BLAST analyses of all protein sequences surrounding the RIP genes yielded maximum scores with arthropod annotated proteins (**<u>Supplementary Table 1</u>**). In contrast, the BtRIPs showed maximum sequence identity to plant RIPs, and marginal identity to their mosquitos’ homologs (around 24%).

**Figure 1.**
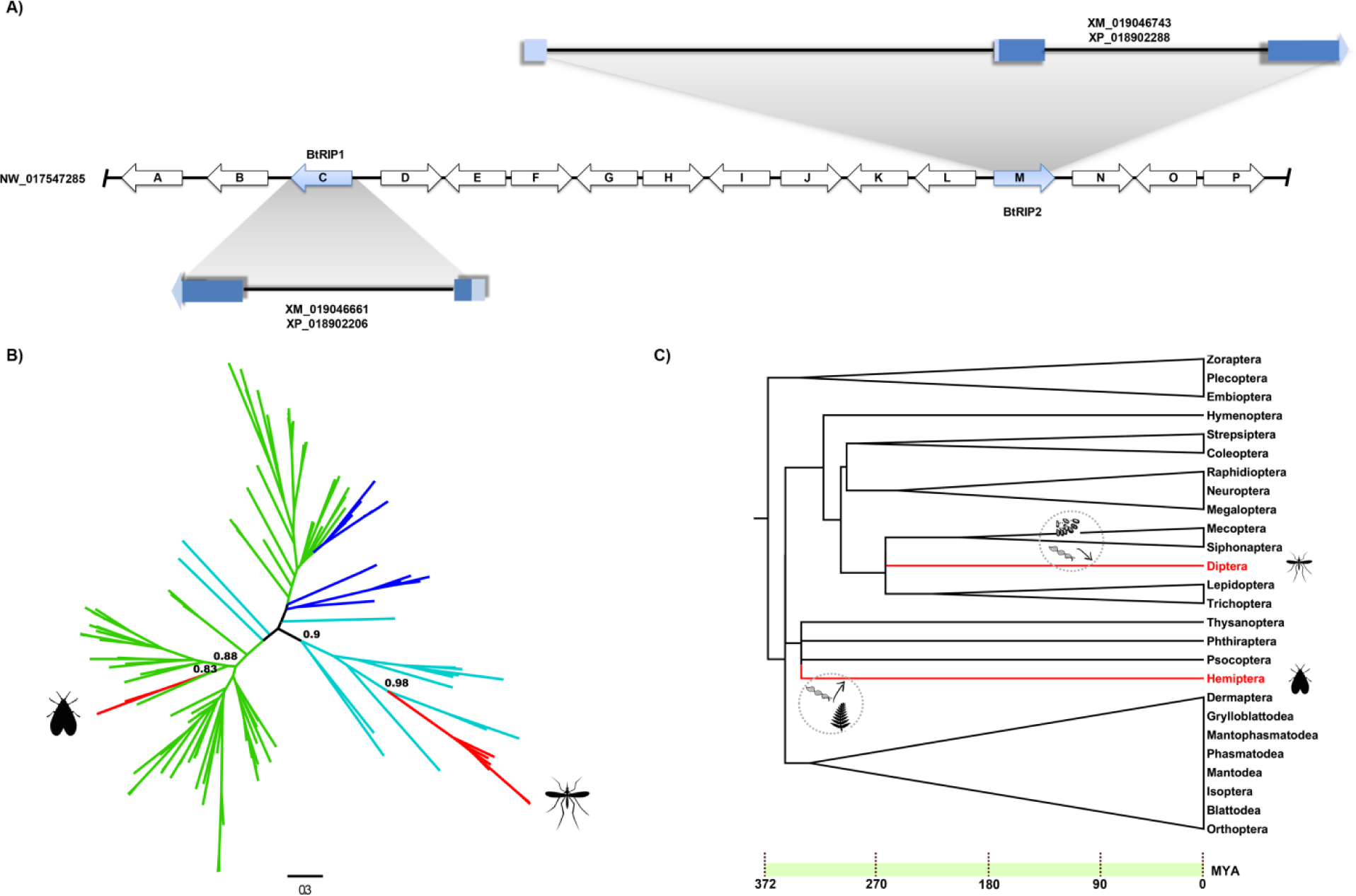
A. Scheme depicting the *B. tabaci* MEAM1 genomic region harboring BtRIP1 (XM_019046661) and BtRIP2 (XM_019046743) genes. Arrows depict the coded proteins in the genomic scaffold. The empty arrows show the surrounding proteins to BtRIPs with highest blast score to arthropod genes (see arrow code in **Supplementary Table 1**). The untranslated (UTRs) and coding regions of mRNA in BtRIP1 and BtRIP2 genes are represented with light blue and blue colors, respectively. **B. Unrooted phylogeny of RIP genes.** Branches are colored according to taxonomy: bacteria (cyan), plants (green), fungi (blue), metazoan (red). TBE support values of major divergences are shown above the nodes. Mosquito and whitefly clades are marked with silhouettes. Fully annotated phylogeny is available at the supplementary material. **C. Neoptera phylogeny.** Tree was obtained from the Time tree knowledge base [28]. Time (MYA) is indicated at the bottom. Diptera and Hemiptera orders are shown in red branches. The two independent HGT events are graphically represented at the estimated times, close to the corresponding branches with hypothetical donors as silhouettes.

We further confirmed the existence of these *Bemisia* newly discovered genes by analyzing two additional, independent genome contigs belonging to the *B. tabaci* SSA-ECA (PGTP01000858; [21]) and *B. tabaci* MED/Q (ML134445; [22]) assemblies. **Supplementary Figure 1** shows sinteny analysis of the genomic regions including RIP genes in these three subspecies of *B. tabaci*, ruling out possible artifacts arisen from sequence assembling. **<u>Supplementary Table 1</u>** summarizes this information.

The BtRIP1 gene is 3466 pb and includes two exons of 273 bp and 517 bp and a single intron of 2730 bp. The mRNA (790 bp) shows 5’ and 3’UTRs of 92 bp and 86 bp, respectively. The mature mRNA encodes a protein of 185 amino acids (**Figure 1A**). The BtRIP2 gene harbors 5974 bp; three exons (156 bp, 354 bp and 563 bp) and two introns (3156 bp and 1745 bp). The 5’ UTR (186 bp) is formed by the whole first exon and 30 bp of the second exon (Figure 1A). The 3’UTR has 65 bp of the last exon. The mature mRNA encodes a protein of 273 amino acids. **<u>Supplementary Table 2</u>** summarizes the features of BtRIPs compared to the average values for the complete genome of *B. tabaci*. Sequence alignment and structural modeling of *B. tabaci* predicted proteins revealed that all the characterized residues forming the active site are conserved. Interestingly, BtRIP1 is remarkably smaller than average RIPs. This is caused by an N-terminal shortening. The functional relevance of this shortening is not clear from the structural model since though a big open cavity is generated, active site architecture is conserved. Other less important *N*-terminal deletions are observed in functional RIPs as lychinin (PDB 2G5X). BtRIP2 is predicted to conserve the architecture and all secondary structure elements canonical of functional RIPs as momordin (PDB 5CF9) or trichosantin (PDB 1QD2) (**Supplementary Figure 2**). By using transcriptomic data (kindly provided by Dr. Zhangjun Fei [19]), we analyzed the expression level of BtRIPs in comparison with the full set of genes. Both genes are expressed at significant level as it can be observed in **Supplementary Figure 3**. In particular, BtRIP2 expression level is at the top quartile of *B. tabaci* genes, strongly suggesting a relevant functional role for this protein.

### 2. BtRIP genes are phylogenetically closer to plant than to mosquito homologues

Next, we performed phylogenetic inferences using the recently discovered BtRIPs along with a representative dataset of RIP sequences from our previous works [13, 14]. The phylogenetic relationships among BtRIPs and previously characterized homologues are expected to shed light on the evolutionary origin and history of these genes and their possible relationship with homologous genes from dipterans. Notably, the tree showed that BtRIPs are embedded in a clade of plant sequences (TBE= 0.96). Interestingly, these RIPs are only marginally related to mosquitoes RIPs, which on the contrary are gathered in a clade of bacterial homologues (TBE= 0.94) (**Figure 1B, Supplementary Figure 4**).

### 3. BtRIPs are derived from a plant genome *via* a single HGT event

To disclose the presence/absence of RIPs in other insects, we performed homology searches using BtRIPs as queries, in complete genome of insects. No hits were retrieved even in other hemiptera species. This fact, along with the phylogenetic inferences, strongly suggests that these genes are not derived from vertical heritage through the insect lineage. Instead, they come from an independent HGT event of a plant RIP gene to an insect genome. **Figure 1C** shows the phylogeny of Neoptera infraclass. As it can be observed, only two clades harbor RIP genes: the lineage including Culicini and Aedini *tribes*, and the lineage that include *B. tabaci* complex species.

As previously reported, Diptera RIP genes are derived from a single HGT event which took place in a window time between the divergence of *Anopheles* and *Culex/Aedes* lineages (around 190 MYA) and before the separation of *Aedes* and *Culex* genus (150 MYA) (Figure 1C). In a similar way, *B. tabaci* RIP genes seem to have been also acquired by HGT, but from a eukaryotic, plant genome source.

The absence of RIP genes in other hemiptera species, strongly suggests that HGT took place in the lineage of Aleyrodidae family (in some time during the last 300 MYA) [30]. The lack of complete genomic data for species more closely related to *B. tabaci* prevents of knowing with more precision the time of the transfer.

We have previously postulated that HGT from bacteria to mosquitoes could have been facilitated by the weakness of the Weisman barrier (the physical separation between somatic and germinal cells in multicellular organisms) at early developmental stages. Feeding of mosquitos’ larva with bacterial species in their ecological niches is consistent with the acquisition of a RIP gene from prokaryotes [14]. Remarkably, findings presented in this work seem to support this hypothesis. At early stages, whiteflies feed from plant sap, and the most plausible source of these horizontally acquired genes is plant genomes. Therefore, our previous and current findings pinpoint that in addition to phylogenetic inconsistencies and taxonomic distribution, developmental and ecological features of animal species should be carefully analyzed when investigating possible HGT events.

## CONCLUSIONS

1. There are two canonical RIP genes in the genome of *B. tabaci*, being the secondly discovered metazoan lineage to harbor genes from this family.
2. *B. tabaci* RIP genes have introns, are transcribed and processed.
3. BtRIP genes are phylogenetically closely related to plant genes, and display only marginal identity to mosquitoes RIP genes.
4. *B. tabaci* RIP genes were acquired by HGT from plants sometime in the last 300 MY.

## Supplementary information

**Figure S1.**
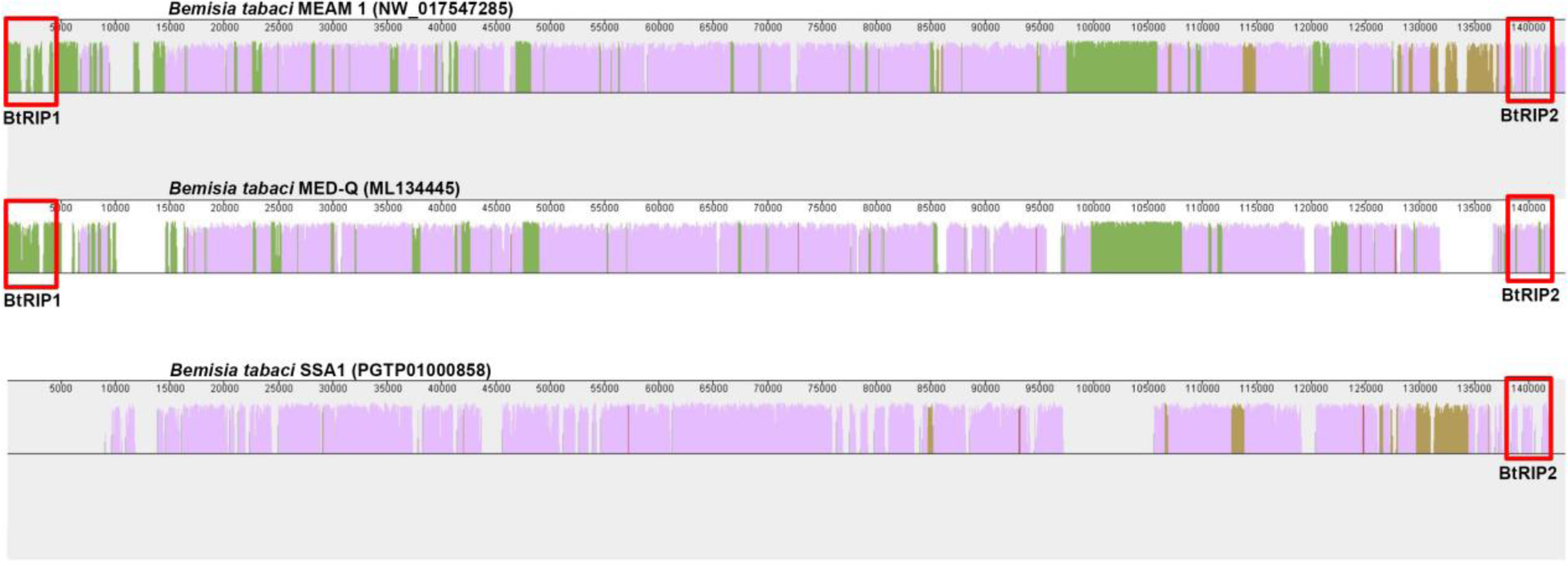
Scheme depicting the *B. tabaci* genomic region harboring BtRIP1 and BtRIP2 genes. The contig region containing RIP genes from *B. tabaci* MEAM 1 (NW017547285), *B. tabaci* MED/Q (ML134445) and *B. tabaci* SSA-ECA (PGTP01000858) were compared using MAUVE software [1]. Sections of the similarity plot colored in violet are conserved among all three genomes; portions in green are conserved among the MEAM 1 and MED-Q sequences, segments conserved among MEAM 1 and SSA 1 are colored in brown. Regions corresponding to BtRIP1 and BtRIP2 are indicated with red boxes.

**Figure S2.**
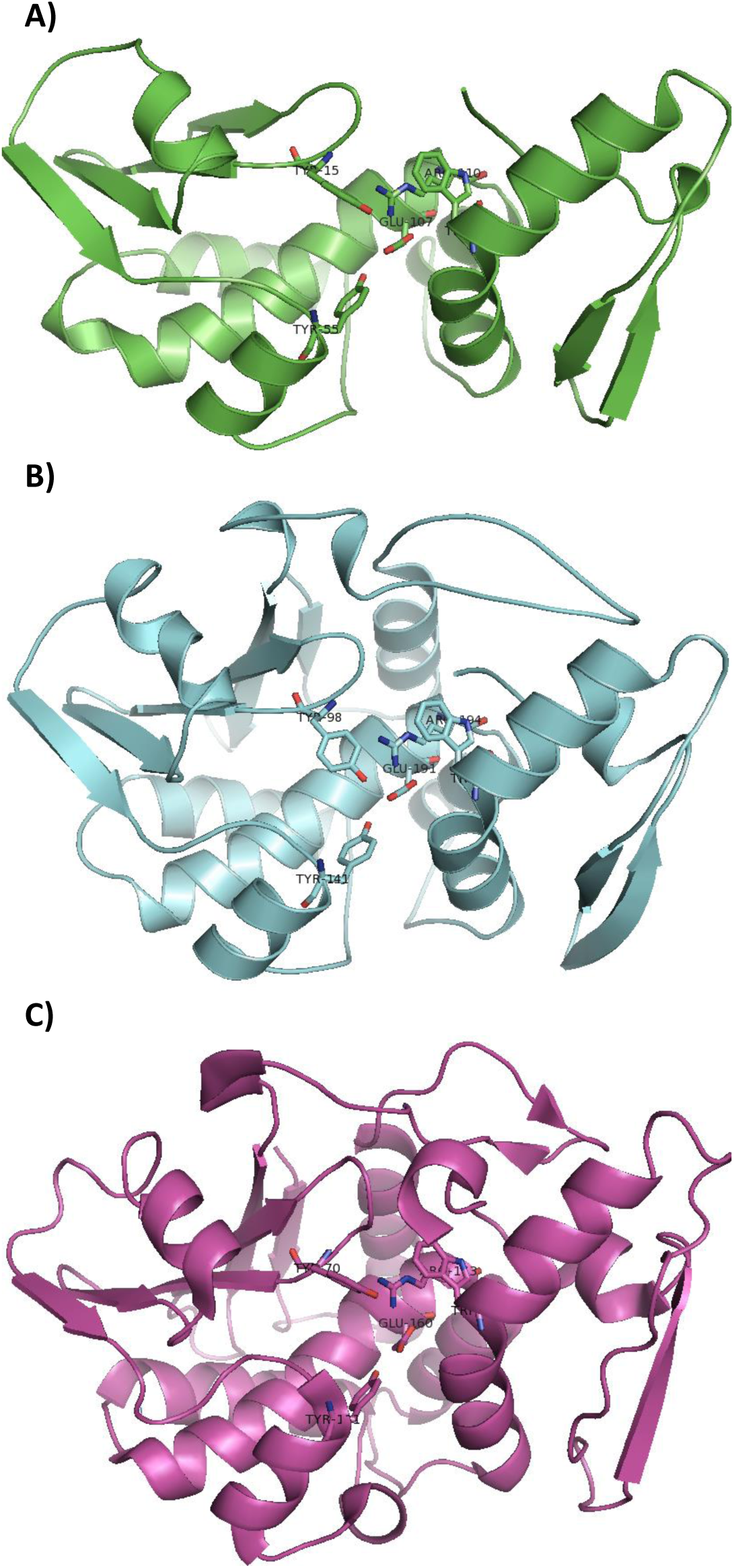
Homology models of BtRIP1 and BtRIP2. Models were constructed in Swiss-Model server [2]. Structures of momordin (PDB 3MY6 or 4YP2) displaying 30-32% sequence identity were used as templates for both BtRIP1 and BtRIP2. **A.** Homology model of BtRIP1 (green). **B.** Homology model of BtRIP2 (cyan) **C.** Structure of momordin PDB 3MY6 (magenta). Active site residues (Y, Y, E, R, W) are numbered and shown as sticks.

**Figure S3.**
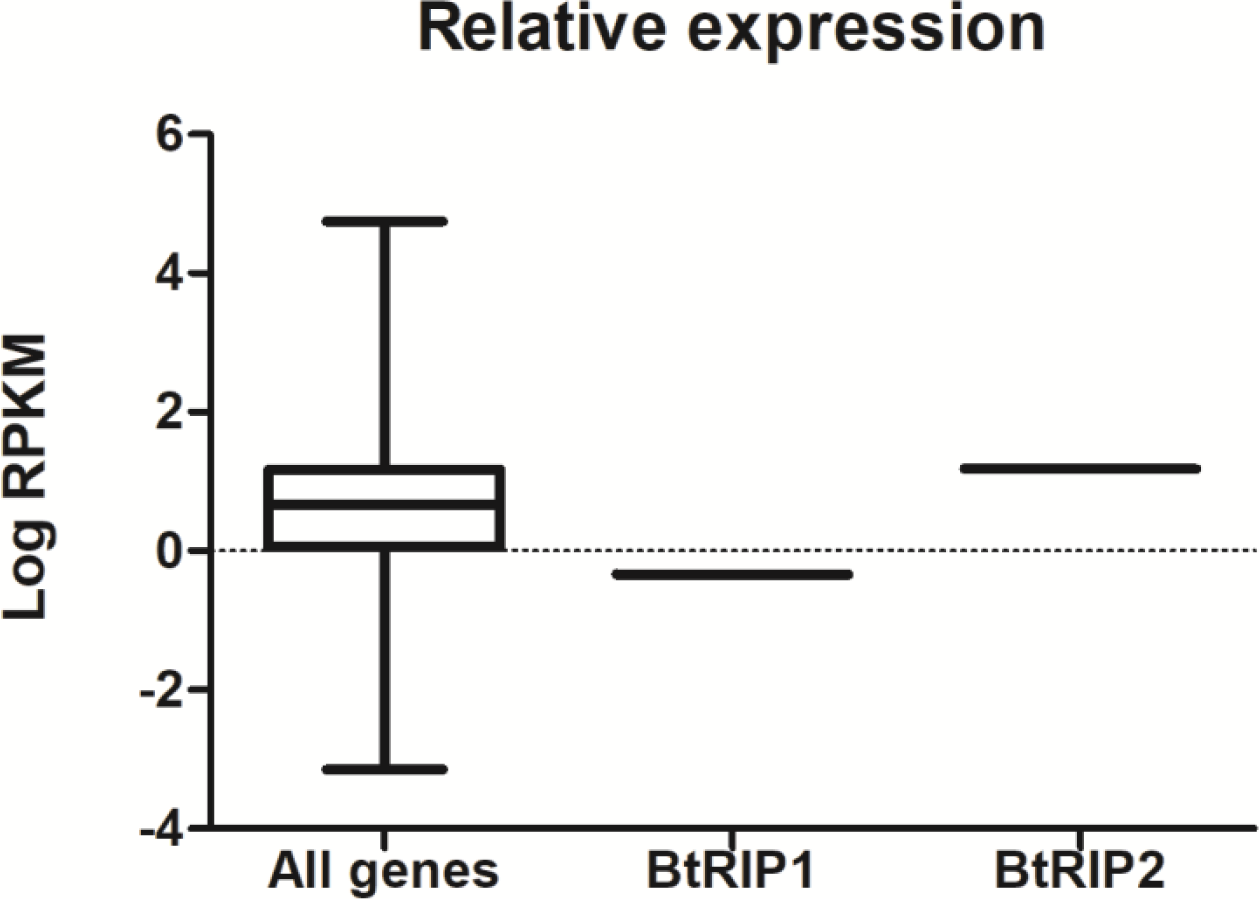
Expression level of BtRIPs. Data provided by Dr. Dr. Zhangjun Fei were plotted as log of RPKM. Values of BtRIP1 and BtRIP2 were compared with the whole set of *B. tabaci* genes according to Chen *et al*. [3]. The box-and-whisker plot shows expression levels as quartiles.

**Figure S4.**
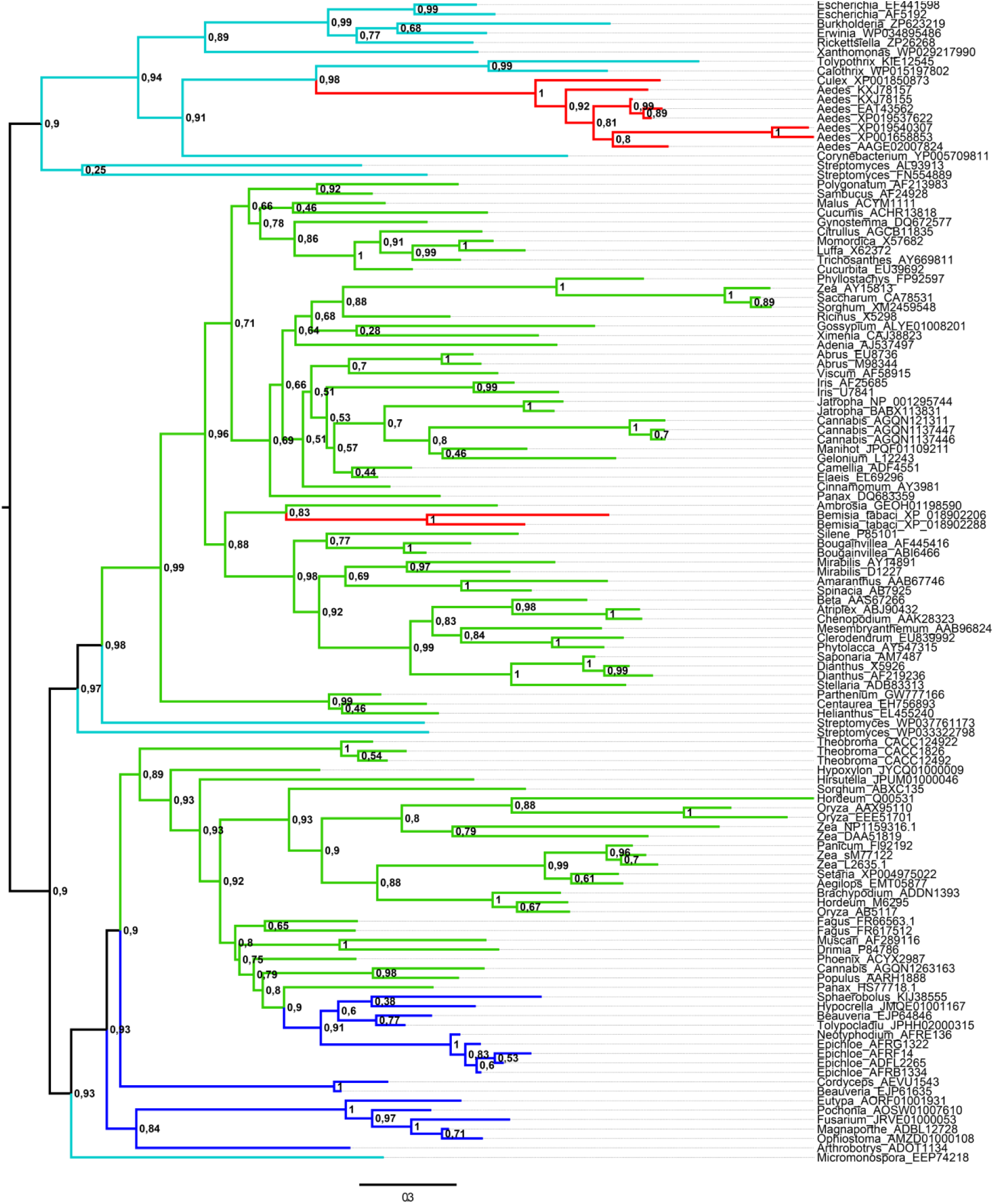
RIPs fully annotated phylogeny. Midpoint rooted phylogeny of RIP genes. Tree was constructed in RAXML 8.2.12. Branches are colored according to taxonomy: bacteria (cyan), green (plants), fungi (blue), metazoan (red). 100 rapid bootstrapping was performed and then transformed to TBE in BOOSTER. TBE support values are shown in nodes. Genus and GenBank accession codes are given for each taxa.

